# Astrocytes Control Recent and Remote Memory Strength by Affecting the Recruitment of the CA1→ACC Projection to Engrams

**DOI:** 10.1101/2023.10.08.561396

**Authors:** Ron Refaeli, Tirzah Kreisel, Maya Groysman, Inbal Goshen

## Abstract

Recent and remote memories are encoded throughout the brain in ‘Engrams’: cell ensembles formed during acquisition, and upon their reactivation, a specific memory can be recalled. The maturation of engrams from recent to remote time points involves the recruitment of CA1 neurons projecting to the anterior cingulate cortex (CA1→ACC). Various modifications to CA1 astrocytes, to the Gq- or Gi-GPCR pathways, during memory acquisition were shown to affect recent and remote recall in seemingly contradictory ways. To address the inconsistency, we applied transgenic approaches for ensemble identification, CLARITY, retro-AAV virus for circuit mapping, and chemogenetics in astrocytes for functional investigation. We manipulated the activity of either Gq- or Gi-pathways in CA1 astrocytes during memory acquisition and tagged cFos+ engram cells and CA1→ACC cells during recent and remote recall in the same animals. The behavioral results were coupled with changes in the recruitment of CA1→ACC projection cells to the engram, demonstrated by the number of CA1→ACC projecting cells in the engram as well as the number of axons projecting from the CA1 engram toward the ACC: Gq pathway activation in astrocytes caused enhancement of recent recall alone and was accompanied by earlier recruitment of CA1→ACC projecting cells to the engram. In contrast, when activating the Gi pathway in astrocytes during acquisition, only remote recall was impaired, and CA1→ACC projecting cells were not recruited during remote memory. Finally, we provide a simple working model, hypothesizing that astrocytes control behavioral performance by targeting the CA1→ACC projection. Specifically, that Gq- and Gi-pathway activation affect memory differently, but do so by modulating the same mechanism. These findings illuminate the importance of astrocytes in the acquisition of fear memory and their implications on recent and remote recall.

## Introduction

Recent hippocampal memory consolidation occurs on the synaptic level during the first hours following learning. Remote memories are consolidated over a much longer time through loosely-defined processes involving multiple brain regions that change to create the persistence of the memory^1,2^, termed together ‘systems consolidation’. For example, prominent theories suggest that the consolidation of remote memories requires hippocampal activation during recent memory, and relies on frontal cortical regions such as the anterior cingulate cortex (ACC) in addition to the hippocampus during more advanced stages^2-15^.

Astrocytes can sense and modify synaptic activity as an integral part of the ‘tripartite synapse’^16-18^, and their role in memory was recently demonstrated repeatedly. Specifically, modification of exogenous GPCRs in astrocytes was used to alter their activity and to examine their real-time involvement in memory. The integration of chemogenetic and optogenetic tools allows reversible manipulation of these cells at the population level: in chemogenetics^19^, intracellular signaling in a genetically defined population is controlled by expressing a designer receptor (most commonly hM3Dq that recruits the Gq-coupled and hM4Di recruiting the Gi-coupled pathways), engineered not to respond to innate ligands, but rather to synthetic, inert drugs like clozapine-N-oxide (CNO)^19^. Optogenetics activates the Gq-coupled pathway with light, e.g. by Opto-α1AR and melanopsin^20,21^.

In general, it can be said that activation of the Gq pathway in astrocytes improves normal memory^20-23^ and enhances it when it is impaired^24^, whereas Gq pathway attenuation blocks memory^25^. On the other hand, activating the Gi pathway in CA1 astrocytes has a harmful effect on memory^26-28^ and blocks beneficial effects on impaired memory^29^. Specially, the results demonstrating the role of astrocytes in recent and remote fear conditioning (FC) memory acquisition, seem contradictory^30,31^: On one hand, hM3Dq, Opto-α1AR and melanopsin activated during acquisition enhance recent memory^20,21^, but on the other, hM4Di activated at the same time does not affect recent memory at all, but impairs remote memory^27^. The aim of this project was to decipher the seemingly opposing effects on memory caused by Gq- and Gi-pathway activation.

When a memory is acquired, a group of neurons with high activity levels are allocated to support it, and some of these neurons will later participate in the recent and remote recall of this memory^32-35^. We recently studied the evolution of CA1 engram cell populations from recent to remote memory recall for the same memory, and found that the populations are stable and that the new cells that join the remote engram are not added randomly during maturation, but rather according to their input and output connections^36^. Particularly, the anterograde CA1 engram neurons projecting to the ACC (CA1→ACC) grow larger in remote memory^36^, and inhibiting CA1→ACC neurons during memory acquisition harms remote recall^27^. Interestingly, Gi pathway activation in astrocytes was shown by us and others to have projection specific effects on dCA1→ACC neurons^26,27^.

Thus, in this study, we manipulated the activity of either Gq or Gi pathway in CA1 astrocytes during contextual FC acquisition and tested the effects on engram cells during recent and remote recall in the same animals, specifically examining the recruitment of CA1→ACC neurons to the engram. Like before^20^, we found that activation of the Gq pathway causes enhancement of recent but not remote recall, while activation of the Gi pathway impairs remote recall performance but spares recent recall. These behavioral results were coupled with changes in recent and remote engram stability, i.e. increased by Gq- and decreased by Gi-pathway activation. Notably, Gq pathway activation on CA1 astrocytes results in early recruitment of the CA1→ACC projection, whereas Gi pathway activation prevents the recruitment of this projection at remote time points. Finally, we suggest a simple working model that synthesizes all of the results, hypothesizing that astrocytes control the behavioral performance by specifically targeting the same mechanism, CA1→ACC projection.

## Results

### Activation of the Gq pathway in CA1 astrocytes during memory acquisition enhances recent memory and strengthens CA1→ACC connectivity simultaneously

Previous studies demonstrated that activation of the Gq pathway in CA1 astrocytes during acquisition of a memory enhances recent recall^20-22^, but its role in remote recall is still unknown. Astrocytes were also shown to affect specific projections in several brain regions^27,37,38^. In a recent paper, we found that the recruitment of the CA1→ACC projection from the CA1 engram increased over time from recent to remote^36^, therefore we wanted to examine whether activating the astrocytic Gq pathway (with hM3Dq) will affect this process. To tag memory engrams, we injected AAV5-cFos-CreER, inducing the expression of Cre under the promoter of the immediate early gene cFos^39-41^ (that is elevated in active neurons) to the dorsal-CA1 (dCA1) and ACC of Ai14-reporter mice (129S6-Gt(ROSA) 26Sor^tm14(CAG-tdTomato)Hze^; see methods), conditionally expressing tdTomato in cells that express Cre. The activity of Cre on tdTomato is limited to a 4-8hr time window^42^ defined by injection of 4-hydroxytamoxifen (4-OHT) (25 mg/kg, intraperitoneal, i.p.), allowing Cre translocation into the nucleus and consequently tdTomato expression. We also injected AAV1-GFAP::hM3Dq(Gq)-mCherry into the CA1, causing expression of hM3Dq in astrocytes, and AAVretro-CaMKII::eGFP into the ACC, tagging the CA1→ACC projection neurons (Figure 1A,B). 4-OHT was administered during recent recall to tag the first engram, and the remote engram was stained in far-red (Alexa Fluor 647) by IHC against cFos (Figure 1B). As astrocytes expressing hM3Dq-mCherry and recent recall engram cells expressing tdTomato are both red, we stained the astrocytes using an α-mCherry antibody and found that >99% of astrocytes were stained yet only <8% of the cfos positive neurons were stained (Figure S1A-C). Animals were trained in contextual FC, and Saline or CNO (3mg/kg, i.p.) were injected 30 minutes before acquisition. As expected^20^, no behavioral differences were observed during acquisition, but memory performance during recent recall was elevated among the mice treated two days prior with CNO compared to the Saline treated controls (F_(5,39)_=30.5, p=1.69*10^-12^, post hoc Saline Recent vs CNO Recent, p=0.049)(Figure 1C). During remote recall, no significant difference between the groups was observed (p>0.42).

**Figure 1:**
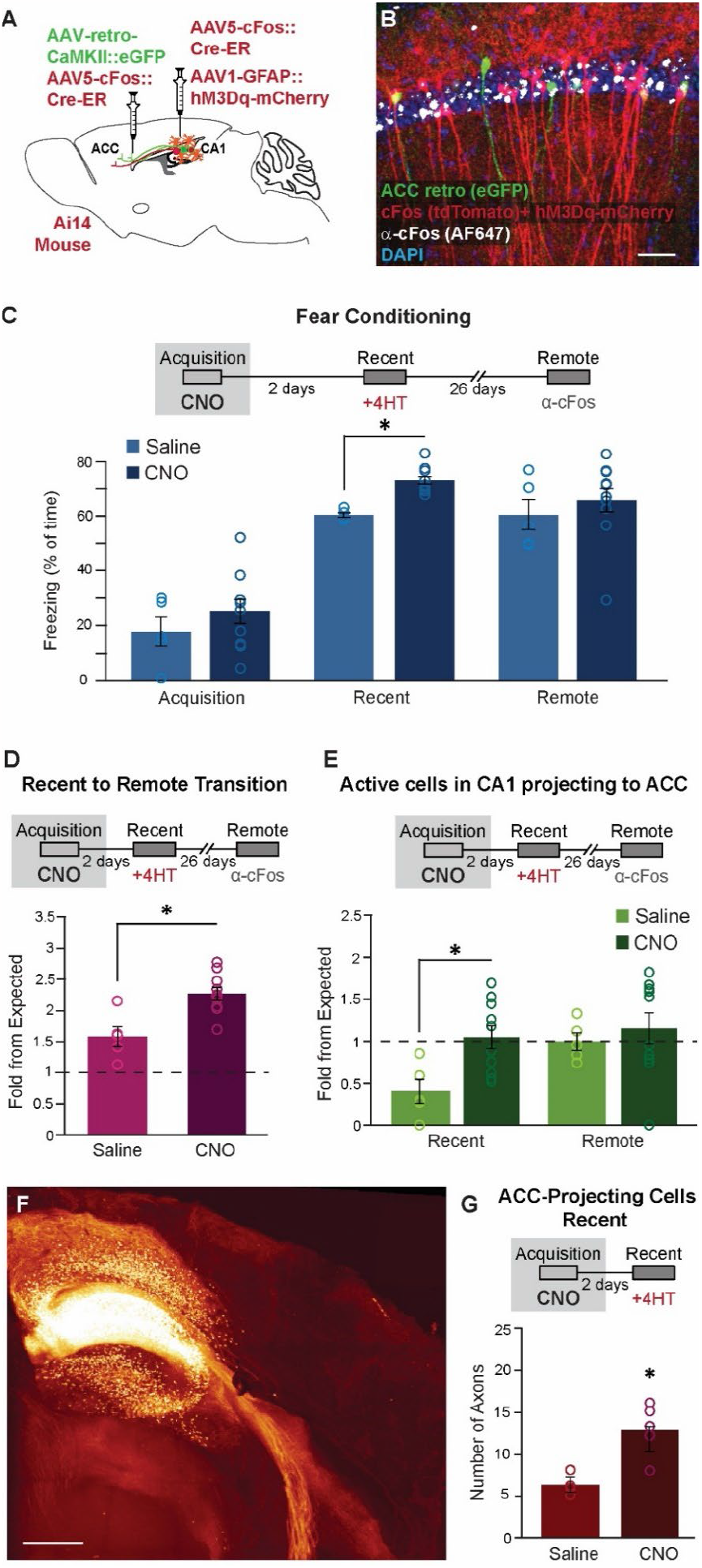
Gq-pathway activation in CA1 astrocytes during memory acquisition enhances recent memory and alters CA1→ACC connectivity. (A) Ai14 mice were injected with a AAV5-cFos::CreER to tag the active cell engram and AAV1-GFAP::hM3Dq-mCherry express hM3Dq in CA1 astrocytes. AAVretro-CaMKII::eGFP was injected into the ACC to tag cells in CA1 projecting to the ACC (CA1→ACC), and AAV5-cFos::CreER to tag the engram in this region. **(B)** CA1→ACC cells express eGFP (green), hM3Dq positive astrocytes express mCherry (red), recent recall cFos positive cells express tdTomato (red), and remote recall cFos positive cells stained with α-cFos (white). Scale bar = 50μm. **(C)** Mice were administered Saline (n=5) or CNO (n=10) during acquisition. Memory enhancement was observed during recent recall (F_(5,39)_=30.5, p=1.69*10^-12^. Post hoc Recent Saline vs. Recent CNO p=0.049) but not during remote recall (Post hoc Remote Saline vs. Remote CNO p=0.53). **(D)** Level of engram reactivation exceeded expected levels for saline (p=0.031) and CNO groups (p=0.000098). The CNO group presented significant higher levels of overlap compared to Saline (t_(13)_=3.74, p=0.0025). **(E)** CNO, compared to the Saline control group, causes significant recruitment of the CA1→ACC neurons during recent, but not during remote recall. (F_(3,26)_=3.26, p=0.038, p recent saline vs. recent CNO = 0.016). **(F)** An entire transparent hemisphere of a mouse expressing tdTomato in cFos positive cells, and hM3Dq-mCherry in astrocytes. The long-range projections of the engram are traceable. Scale bar = 1mm. **(G)** The number of tdTomato positive axons of ensemble cells heading toward the ACC significantly increased when CNO (n=4) was introduced to mice expressing hM3Dq in CA1 astrocytes during memory acquisition, compared to the Saline control group (n=5; p=0.0058). Data presented as mean ± SEM.

By tagging the engram cells during recent and remote memory within the same mouse, we investigated whether the reactivation levels changed between the Saline and the CNO treated groups. We found that both groups exceeded the expected reactivation level (t-test t_(4)_=3.27, p=0.0031 and t_(9)_=6.61, p=0.000098, respectively), but remarkably, the similarity between the recent and remote engram in the CNO treated group was significantly higher (t_(13)_=3.74, p=0.0025)(Figure 1D). In the same mice, the reactivation level was measured at the ACC as well, and while both groups exceeded the expected level of reactivation (t_(4)_=5.58, p=0.005 for Saline and t_(9)_=6.43, p=0.00012 for CNO), no effect of CNO was found between the groups (Figure S1D).

Studying the contrasts between recent and remote memory engrams demands an understanding of the dynamics within the long-range connections between areas of the brain. While the connections between any two brain regions remain constant, the selected cells that comprise the engrams within these regions can alter based on the target of their projection. By examining the anterograde connections of hippocampal engram neurons at various time points, we can uncover the evolving relations with their downstream brain structures. In a previous work, we showed that one difference between recent and remote CA1 engrams is that the later engram recruits more CA1→ACC projection cells^36^. We sought to determine whether astrocytes in which the Gq pathway was activated during acquisition can affect this process. The ratio of these projection cells among engram cells could be measured by injecting mice with AAVretro-CaMKII::eGFP to ACC which labels all CA1→ACC cells in green, in addition to AAV5-cFos-CreER. In the Saline control group we saw, as before^36^, a significant increase in the likelihood of the CA1→ACC projection to participate in the remote engram compared to the recent engram (F_(3,26)_=3.26, p=0.038; post hoc Saline recent vs Saline remote, p=0.049), with the recent recall measures below the expected values^36^. Extraordinarily, in the CNO group, the likelihood of the CA1→ACC group of cells in the engram was already elevated during recent recall (post hoc of Recent CNO vs Recent Saline, p=0.016)(Figure 1E) to values similar to those usually found during remote (p=0.6).

We next demonstrated the early recruitment of the CA1→ACC projection following astrocytic Gq pathway activation during acquisition by using another technique: we imaged and counted the number of engram axon projections from the dCA1 in whole brains that underwent CLARITY^42-44^. We tagged the active recent recall cells as we did before: Ai14 mice were injected with cFos::Cre-ER and AAV1-GFAP::hM3Dq(Gq)-mCherry into the dCA1 (Figure 1A) and administered CNO 30 minutes before FC acquisition and then 4-OHT one hour after behavior. Since behavioral changes after Gq activation in astrocytes occur only during recent recall, we targeted this time point. Again, memory performance during recent recall was elevated among the mice that were treated with CNO compared to the Saline treated controls (F_(3,14)_=90.6, p=2.075*10^-9^, post hoc Recent CNO vs Recent Saline, p=0.001)(Figure S1E). Four weeks after tagging the active cells, mice were sacrificed and a CLARITY procedure was performed in order to turn the brains transparent, allowing imaging of full hemispheres with single axon resolution (Figure 1F). The dCA1 projects to several targets, three of them, in descending order, are the mammillary bodies (MB), the nucleus accumbens (NAc), and the ACC. For both groups of mice, we found the same total number of axons projecting from the dCA1 engram cells (p>0.9, Figure S1F), and no significant changes in the number of axons between the groups in the MB (p=0.25)(Figure S1G) or the NAc (p=0.62)(Figure S1H) were found. We did however find that the CNO treated mice, i.e. those with recruited Gq pathway in astrocytes during acquisition, showed a higher number of CA1 axons projecting to the ACC compared to the Saline injected group during recent recall (t-test; t_(7)_=3.9, p=0.0058)(Figure 1G).

These results show that activation of the Gq pathway in CA1 astrocytes during acquisition expedites CA1→ACC recruitment during recent recall, seen first by the quantity of these projection cells in the engram, and second by the elevated number of engram cell axons projecting to the ACC from dCA1.

### Activation of the Gi pathway in CA1 astrocytes during acquisition impairs remote memory and degrades the recruitment of the CA1→ACC projection.

The effect of Gi pathway activation in astrocytes on memory was examined by several groups and found to be detrimental^26-29^. Specifically, it was shown that Gi pathway activation in CA1 astrocytes during memory acquisition impairs remote recall, however it does not affect recent recall^27^. We wanted to test whether Gi pathway activation in astrocytes also has an effect on CA1 engrams and on the CA1→ACC projection recruitment in them.

To that end, Ai14 mice were injected with AAV5-cFos::CreER and AAV8-GFAP::hM4Di(Gi)-mCherry into the CA1, as well as AAV5-cFos::CreER and AAVretro-CaMKII::eGFP into the ACC (Figure 2A-B). The recent recall engram was tagged by 4-OHT administration allowing the Cre to enter the nuclei of active cells during retrieval thus expressing tdTomato in the Ai14 mice, and the remote engram was stained by IHC against cFos. As before, astrocytes expressing hM4Di-mCherry and recent engram cells expressing tdTomato are both red, so again, we stained for mCherry and found that >95% of astrocytes were stained and only <8.5% of the cfos positive neurons (Figure S2A-B). Animals were trained in contextual FC, and Saline or CNO (10 mg/kg, i.p.) was injected 30min before acquisition. No behavioral difference was observed during acquisition or recent recall, but memory performance during remote recall was reduced among CNO treated mice compared to the Saline controls (F_(5,18)_=75.94, p=1.85*10^-11^, post hoc Remote CNO vs Remote Saline, p=0.0008)(Figure 2C), as we have shown before^27^. When examining the reactivation levels between recent and remote engrams, we found that while the Saline group exceeded the level of expected overlap (t_(3)_=7.22, p=0.0055), as was previously shown^36^, the CNO group did not differ from chance level (t_(3)_=0.93, p=0.422), resulting in a significant difference between the groups (t_(6)_=3.24, p=0.018)(Figure 2D). The level of reactivation was measured at the ACC as well, and here too, the Saline group was significantly higher than the expected reactivation level (t_(3)_=3.19, p=0.049)(Figure S2C), however there was no significant difference between Saline and CNO (p=0.95).

**Figure 2:**
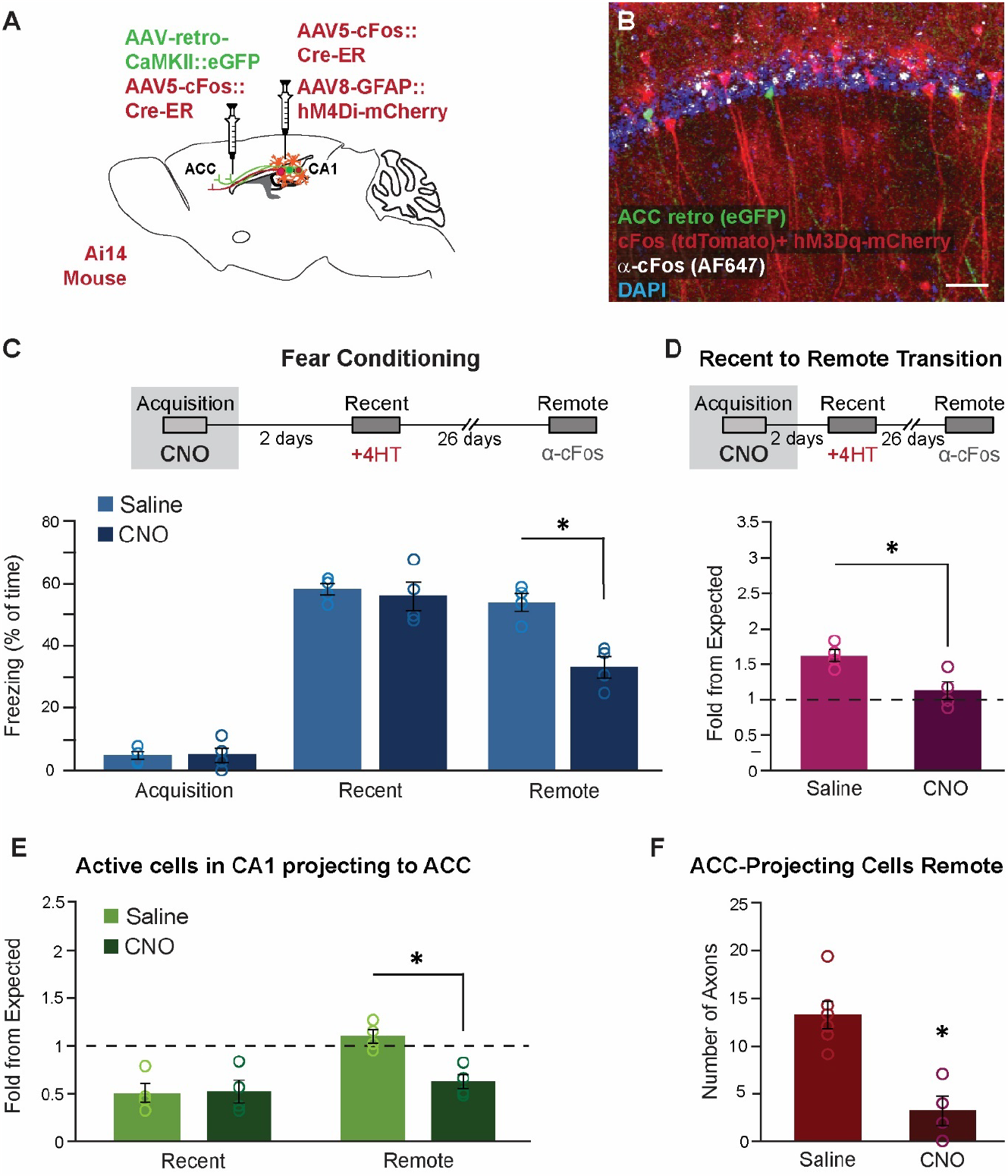
Activation of the Gi pathway in CA1 astrocytes during memory acquisition impairs remote memory and degrades the recruitment of the CA1→ACC projection. (A) Ai14 mice were injected with a AAV5-cFos::CreER and AAV8-GFAP::hM4Di-mCherry into the CA1 to tag active cells and express hM4Di in astrocytes, as well as AAV-retro-CaMKII::eGFP to tag cells in CA1 projecting to the ACC (CA1→ACC) and AAV5-cFos::CreER into the ACC. **(B)** CA1→ACC cells express eGFP (green), hM4Di positive astrocytes express mCherry (red), recent recall cFos positive cells express tdTomato (red), and remote recall cFos positive cells were stained ex-vivo (white). Scale bar = 50μm. **(C)** Mice were administered Saline (n=4) or CNO (n=4) during memory acquisition. Memory impairment was observed during remote recall (F_(5,18)_=75.94, p=1.85*10^-11^, post hoc Remote Saline vs. Remote CNO p=0.0008) but not during recent recall (p=0.996). **(D)** Engram reactivation exceeded expected level for Saline (t_(3)_=7.22, p=0.0055) but not for the CNO group (p=0.42), resulting in a significant difference between the groups (t_(6)_=3.24, p=0.018). **(E)** CNO, compared to the Saline control group, caused a significant reduction in the recruitment of CA1→ACC neurons during remote recall (F_(3,12)_=9.05, p=0.002, post hoc Remote Saline vs. Remote CNO p=0.016) but causes no changes during recent (p=0.999). **(F)** During remote recall, the number of tdTomato positive axons directed toward the ACC decreased significantly when CNO (n=4) was introduced during memory acquisition to mice expressing hM4Di in CA1 astrocytes compared to the Saline control group (n=6)(p=0.0017). Data presented as mean ± SEM.

Next, we tested whether astrocytes in which the Gi pathway was activated during acquisition could affect the recruitment of the CA1→ACC projection to recent and remote CA1 engrams. Recruitment was measured by counting the ratio of green CA1→ACC cells among engram cells. In the Saline control group we saw, as before^36^, a significant increase in likelihood of the CA1→ACC projection to take part in the remote engram compared to the recent engram (F_(3,12)_=9.05, p=0.002; post hoc Saline Recent vs Saline Remote, p=0.003). Remarkably, in the CNO group the likelihood of the CA1→ACC did not increase in remote recall (post hoc CNO Recent vs CNO Remote, p=0.872)(Figure 1E), resulting in a significant difference from the Saline group during remote recall (post hoc of Remote CNO vs Remote Saline, p=0.016).

Lastly, we tested whether failure to recruit the CA1→ACC projection during remote recall following astrocytic manipulation during acquisition is apparent in the number of dCA1 engram cells that send axons to the ACC in the remote engram. To that end, a group of Ai14 mice were injected with the AAV5-cFos::CreER and AAV1-GFAP::hM4Di(Gi)-mCherry to the dCA1 and underwent FC with Saline or CNO injection 30min before acquisition. Since behavioral changes after Gi pathway activation in astrocytes occur solely during remote recall, we targeted this time point. Memory performance during remote recall was diminished among the CNO mice compared to the Saline controls (F_(3,16)_=133, p=1.6*10^-11^)(Figure S2D). Four weeks after tagging the active cells, mice were sacrificed and a CLARITY procedure was performed. For the CNO group of mice, we found a small but significant reduction in the total number of axons projecting from the dCA1 engram cells (t_(8)_=2.86, p=0.021)(Figure S2E). We found no significant changes in the number of axons between the groups in the MB (p=0.45)(Figure S2F) or the NAc (p=0.758)(Figure S2G). Regarding the projections to the ACC, as we recently showed^36^, their number in the Saline group in remote memory is doubled compared to recent (Figure 2F compared to Figure 1F). However, we found that the CNO treated mice, i.e. those with recruited Gi pathway in astrocytes during acquisition, showed a lower number of CA1 axons projecting to the ACC during remote recall compared to the Saline injected group (t-test; t_(8)_=4.64, p=0.0017)(Figure 2F).

The results of this section show that activation of the Gi pathway in CA1 astrocytes during acquisition degrades CA1→ACC recruitment during remote recall, as seen in the number of projection cells in the engram and in the reduced number of engram cell axons going to the ACC from dCA1.

### The conflicting effects of Gq and Gi pathway activation in CA1 astrocytes on memory explained by their contradictory influence on the recruitment of the CA1→ACC projection to recent and remote engrams

The results above together show that at different time points, CA1 astrocytic manipulation can enhance or impair memory and control the CA1→ACC recruitment to the engram. Based on the fact that these effects are caused by astrocytic manipulations at the same time, we suggest the following working model: Activating the Gq pathway in CA1 astrocytes during memory formation promotes CA1→ACC recruitment to the engram at an early time point (Figure 3A*i*) when the recruitment level is normally low, and via this change it enhances recent memory performance (Figure 3B*ii*). At the remote time point, the same Gq pathway manipulation does not cause any change from the normal level of CA1→ACC recruitment, and thus it does not affect memory. Activation of the Gi pathway in CA1 astrocytes during memory acquisition prevents recruitment of the CA1→ACC projection at the remote time point (Figure 3A*iii*) when the recruitment level is normally high, thus causing an impairment in remote memory performance (Figure 3B*iv*). Since there is no effect on CA1→ACC projection recruitment at an early time point, there is also no effect on recent memory performance at that time.

**Figure 3:**
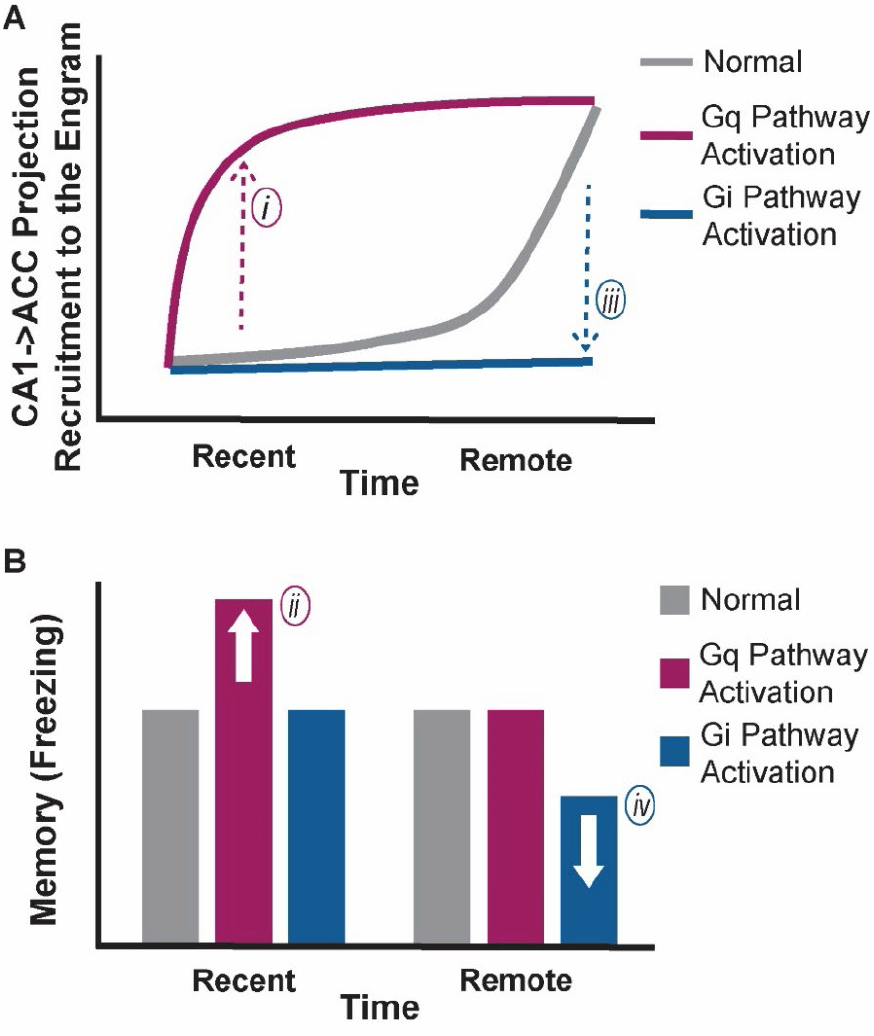
Working model settling the conflicting effects of Gq and Gi pathway activation in CA1 astrocytes on memory. The changes caused by different ways of astrocytic modulation on CA1→ACC recruitment to the engram **(A)** may explain their opposing effects on memory **(B)**.

The results suggest, that the contrasting behavioral result do not stem from two separate mechanisms activated by the Gq- and Gi-pathways, but from opposing effects on a single mechanism – the recruitment of CA1→ACC projection to the engrams.

## Discussion

This study provides the first demonstration of an astrocytic effect on memory engrams, their stability, and their configuration. We manipulated the activity of either the Gq-or the Gi-pathways in CA1 astrocytes during memory acquisition and tagged engram cells during recent and remote recall in the same animals. We found that altering the performance of CA1 astrocytes solely during the formation of the contextual FC causes diverse effects on behavioral performance – activating the Gq pathway causes enhancement of recent recall alone, while activating the Gi pathway impairs remote recall performance but does not affect recent recall. Interestingly, these behavioral results were coupled with changes in CA1→ACC projecting cells recruitment to the engram – demonstrated by the amount of this projection cells in the engram as well as the number axons projecting toward the ACC from dCA1. Higher levels than normal accompanied improved memory, and lower levels than normal coupled with impaired recall.

In a recent study, we found that the CA1 engrams remain stable between recent and remote recall and that inhibition of the engram for recent recall during remote recall functionally impairs memory^36^. We also found that the new cells that joined the remote recall engram during maturation in the CA1 and are not added randomly but differ according to their connections, for instance, the portion of anterograde CA1→ACC engram cell projection grows larger^36^. The current study suggests that these changes in CA1→ACC projection recruitment are important for memory performance, as recruitment increased parallel to enhanced memory and decreased concomitantly with impaired recall.

The processes of engram allocation and maintenance depend on the activity level of neurons^33,45^ which may well be contingent upon their environment. Each astrocyte in the CA1 engulfs approximately 14 pyramidal cell somata^46^, envelopes multiple synapses, and can modify activity^47-50^. In recent years, it has been shown that specific manipulation of astrocytes can modify recent and remote memory: when CA1 astrocytes are manipulated by activating the Gq pathway, recent normal memory is improved^20-23^, impaired memory is enhanced^24^, and if the Gq pathway is attenuated, memory is blocked^25^. Gi pathway activation in astrocytes has the opposite effect - it blocks schema establishment^26^, and whereas its activation during FC acquisition has no effect on recent recall, it dramatically decreases remote recall^27^. Chronic Gi activation attenuates stress-enhanced FC^28^, and cancels the beneficial effect of CA3 activation on short-term memory in APP/PS1 mice^29^. It seems that Gq and Gi have opposite roles in memory. During FC especially, Gq pathway activation enhanced recent memory^20^, and Gi pathway activation impaired remote memory^27^. The latter manipulation was shown to specifically modulate the CA1→ACC projection^27^, indicating that CA1 astrocytes can bear influence on memory at the resolution of a singular projection. Indeed, the current study shows that Gq pathway activation in astrocytes, like the Gi pathway, effects this specific projection during FC acquisition and suggests that the difference stems from the contrary effects these pathways have on CA1→ACC projection recruitment to the CA1 engram of the memory, as indicated by the number of ACC projecting cells in the engram and in the quantity of axons heading toward the ACC.

While the effects of chemogenetic agents in neurons are clear: hM3Dq manipulating the Gq-coupled pathway is excitatory, and hM4Di, manipulating the Gi-coupled pathway, is inhibitory^19^, the picture in astrocytes is more complex^31,51^. A limiting factor is that there is no direct indicator for astrocytic activity beyond intracellular astrocytic calcium (Ca^2+^) elevations, that serve as a proxy to activity, but can signals many other effects caused by various mechanisms. hM3Dq in astrocytes increases Ca^2+^ in these cells^20,52^, so it seems to be excitatory. However, hM4Di also increases Ca^2+^ activity in cortical astrocytes^52-54^ immediately when applied and for some time, but in the hippocampus, it inhibits Ca^2+^ activity on behaviorally relevant time-scales (30min)^27^ (but see^54^). Clearly, the question of how to observe the dissimilar effects of different chemogenetic agents on astrocytes is far from being solved. However, their effect via astrocytes on neuronal activity^20,27,31,53,54^, behavior^20,22,26-28,31,53^, and, in this paper, on engram projection is clear.

These findings illuminate the function of astrocytes during acquisition of fear memory and the implications on recent and remote recall. It demonstrates how astrocytes affect the CA1→ACC projection recruitment to the engram ensemble, in diverse directions by different GPCRs, and we suggest a working model describing the relation between the change in the engram and the difference in memory.

## Supplementary Material

**Figure S1:**
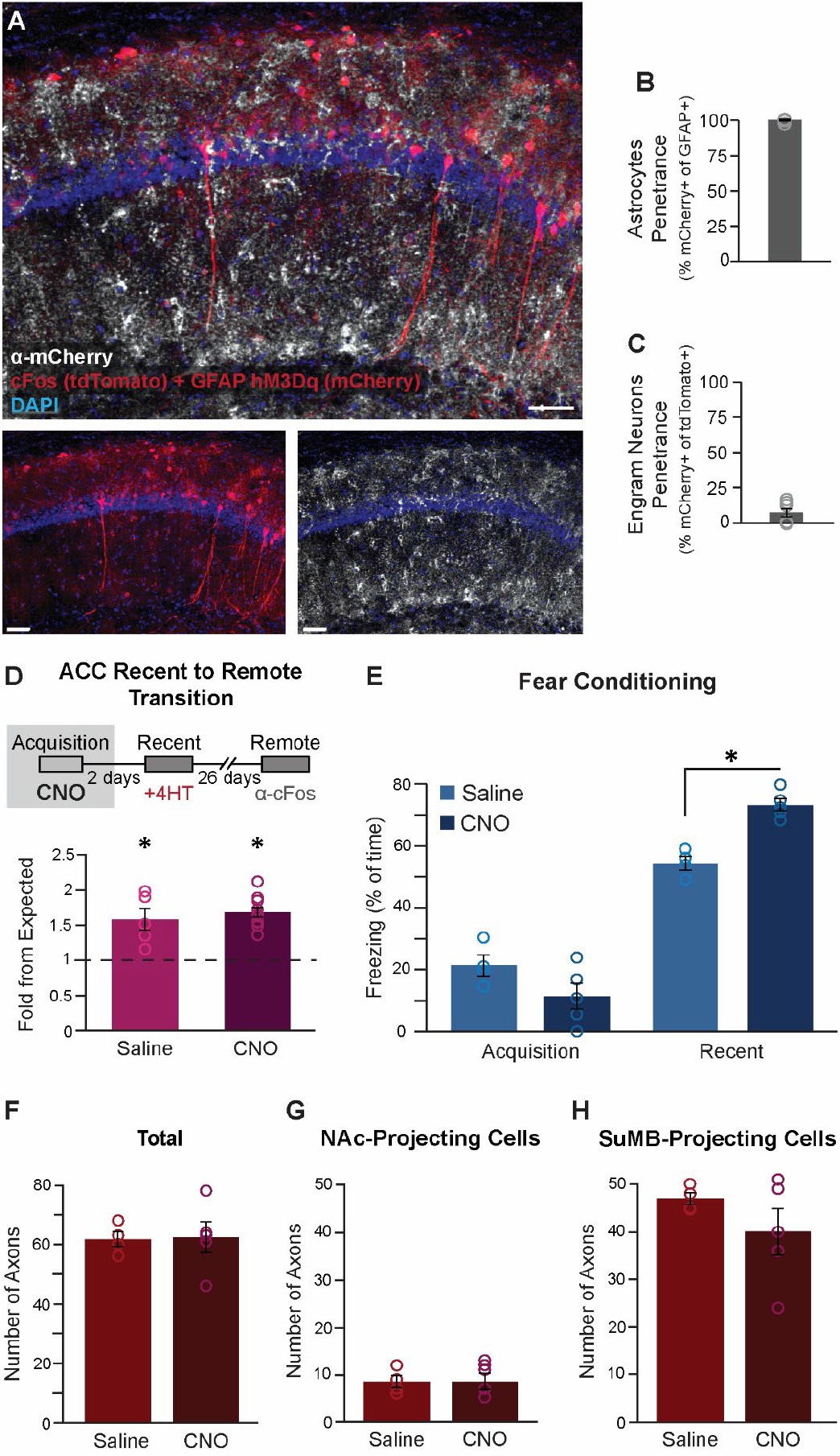
hM3Dq is expressed in astrocytes together with cFos tagged neurons. (**A**) cFos positive cells express tdTomato (red), hM3Dq positive astrocytes express mCherry (red) and α-mCherry staining in AF647 (white). Scale bars = 50μm. mCherry staining reveals penetrance for astrocytes >99% (262 out of 264 astrocytes express mCherry)**(B)** but only >8% for cfos positive neurons (5 out of 105 neurons were positive to mCherry too)**(C). (D)** Level of engram reactivation exceeded expected levels for saline (t_(4)_=5.58, p=0.005) and CNO groups (t_(9)_=6.43, p=0.00012). No significant changes were observed between the groups (p=0.505). **(E)** For the CLARITY experiments, mice were administered saline (n=4) or CNO (n=5) during acquisition. Memory enhancement was observed during recent recall (F_(3,14)_=90.6, p=2.075*10^-9^, post hoc Recent Saline vs. Recent CNO, p=0.004). **(F)** The number of tdTomato positive axons of ensemble cells leaving the dCA1 in cleared brains did not alter between the CNO and the Saline groups. No significant changes were observed when counting the axons heading toward the NAc **(G)** or the MB **(G)** between saline and CNO groups. Data presented as mean ± SEM.

**Figure S2:**
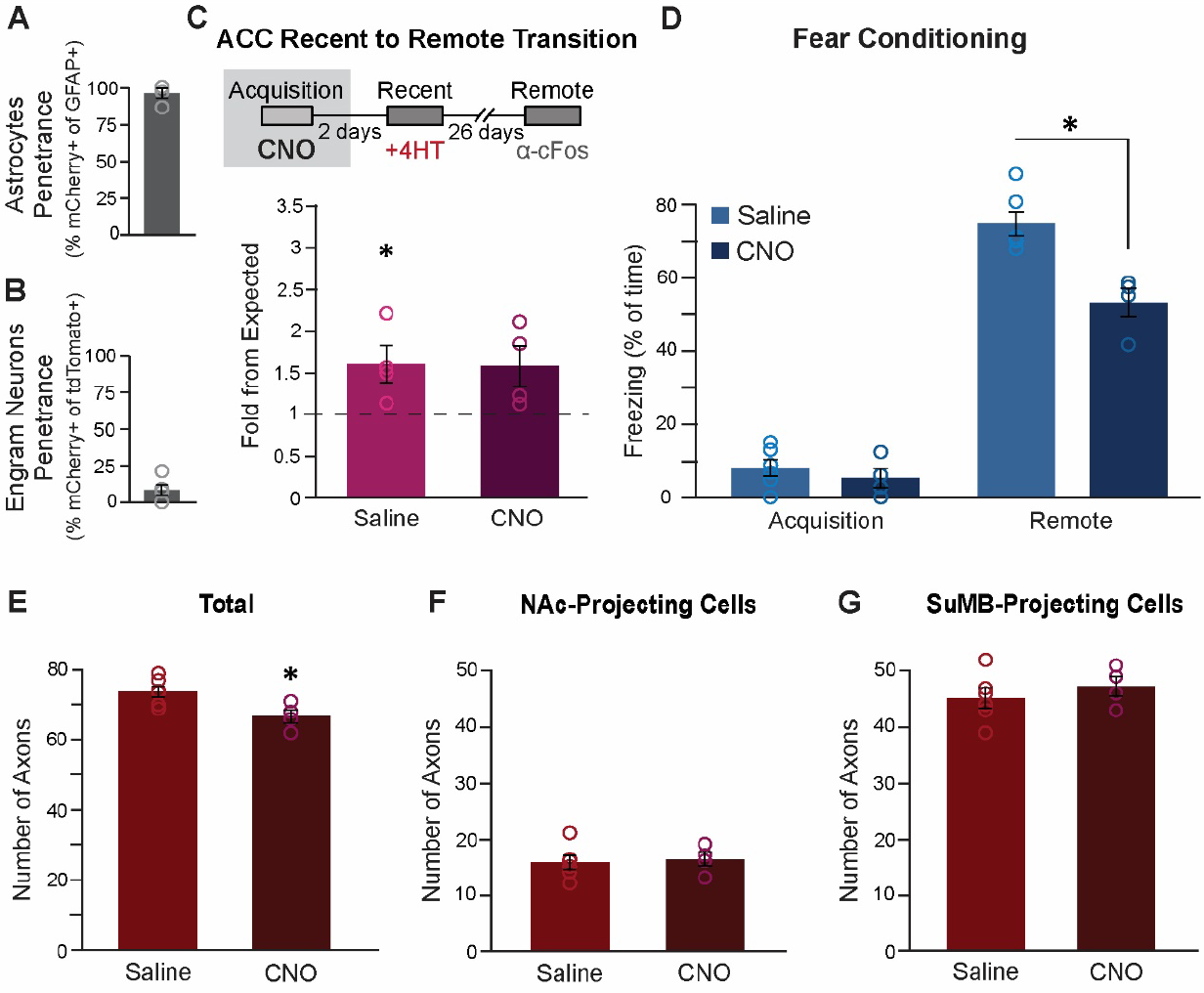
hM4Di is expressed in astrocytes, together with cFos tagged neurons. **(A)** mCherry staining reveals penetrance >95% in astrocytes (113 out of 119 astrocytes express mCherry) **(B)** but >8.5% in cfos positive neurons (9 out of 82 neurons were positive to mCherry too). **(C)** Engram reactivation exceeded the expected levels for saline (p=0.049) but not CNO (p=0.077). No significant changes were observed between the groups (t_(6)_=0.07, p=0.95). **(D)** For the CLARITY experiments, mice were administered saline (n=6) or CNO (n=4) during acquisition. Memory enhancement was observed during remote recall (F_(3,16)_=133, p=1.6*10^-11^, post hoc Remote Saline vs. Remote CNO, p=0.0009). **(E)** The number of tdTomato positive axons of ensemble cells stemming from the dCA1 in cleared brains decreased in the CNO treated mice compare to the Saline treated, but no significant changes were observed when counting the axons heading toward the NAc **(F)** or the MB **(G)** between saline and CNO groups. Data presented as mean ± SEM.

## Methods

### Mice

*Ai14 mice*: B6;129S6-Gt(ROSA)26Sor^tm14(CAG-tdTomato)Hze^, J-Stock No: 007908. These mice contain a loxP cassette with a stop codon, followed by CAG promoter-driven red fluorescent protein (tdTomato)^55^.

*Excluded mice*: Mice were excluded if no viral expression was visible (post mortem).

### Stereotactic Injections

Mice were anesthetized with isoflurane, and their heads were placed in a stereotactic apparatus (Kopf Instruments, USA). The skull was exposed, and a small craniotomy was performed. Mice were bilaterally microinjected using the following coordinates - for the CA1: Anteroposterior (AP), -1.85mm from Bregma, Mediolateral (ML), ±1.4mm, Dorsoventral (DV), -1.5mm. For the ACC: AP +0.35mm, ML ±0.35mm, DV -1.8mm. All microinjections were performed using a 10μL syringe and a 34-gauge metal needle (WPI, Sarasota, USA). The injection volume and flow rate (0.1 ml/min) were controlled by an injection pump (WPI). Following each injection, the needle was left in place for 10 additional minutes to allow for diffusion of the viral vector away from the needle track. The needle was then slowly withdrawn. The incision was closed using sewing and Vetbond tissue adhesive. For postoperative care, mice were subcutaneously injected with Tramadex (5mg/kg).

### Viral Vectors

AAV5-cFos::cre^ER^, AAVretro-CaMKII::GFP, AAV1-GFAP::hM3D(Gq)-mCherry and AAV8-GFAP:: hM4D(Gi)-mCherry were all from the ELSC Vector Core Facility.

### Fear Conditioning

The FC apparatus consisted of a conditioning cage with a grid floor wired to a shock generator and surrounded by an acoustic chamber. To induce FC, mice were placed in the cage for two minutes, and a pure tone (2.9 kHz) was then sounded for 20 seconds followed by a two second foot shock (0.6mA). This procedure was then repeated, and 30 seconds after the delivery of the second shock, mice were extracted from the conditioning cage. Fear was assessed by a continuous measurement of freezing (complete immobility), the dominant behavioral fear response. To test contextual FC, mice were placed in the original conditioning cage, and freezing was measured for five minutes. Contextual FC recall was measured at day 2 and day 28.

### 4-hydroxytamoxifen (4-OHT)

For Ai14 mice, 3 mg of 4HT (Sigma H7904) were dissolved in 120μl of ethanol, and 480μl of sunflower oil were then added to create total volume of 600μl. Each solution was used on the day of preparation. One hour after the relevant memory related task, 4-OHT solution was given to all mice (25mg/kg; i.p.) to enable CreER-mediated recombination.

### CNO

CNO (Tocris #4936) was dissolved in DMSO and then diluted in 0.9% saline to yield a final DMSO concentration of 0.5%. Saline solution for control injections also consisted of 0.5% DMSO. 3mg/kg was intraperitoneally injected 30min before the behavioral assays for the Gq pathway activation^20^, and 10mg/kg CNO similarly for the Gi pathway activation^27^. The chosen doses of CNO did not induce any behavioral signs of seizure activity.

### Immunohistochemistry

90 minutes after the final memory task, mice were transcardially perfused with cold PBS followed by immediate removal of the brain into 4% paraformaldehyde (PFA) in phosphate-buffered saline (PBS). Brains were postfixed overnight at 4#x00B0;C and cryoprotected in 30% sucrose in PBS. Brains were sectioned to a thickness of 50 μm using a sliding freezing microtome (Leica SM 2010R) and preserved in a cryoprotectant solution (25% glycerol and 30% ethylene glycol in PBS). Free-floating sections were washed in PBS and incubated for 1 hr in blocking solution (1% of bovine serum albumin, BSA, and 0.3% Triton X-100 in PBS). For cFos staining, the relevant brain slices were incubated for 7 days at 4#x00B0;C with rabbit anti-cFos primary antibody (1:10000, Synaptic system, #226003), and for the mCherry staining, slices were incubated overnight at 4#x00B0;C with rabbit anti-mCherry primary antibody (1:200, Invitrogen, #PA5-34974). Sections were then washed with PBS and incubated for 2 hr at room temperature with a secondary antibody (1:500, AF 647, donkey anti rabbit, #711-605-152, Jackson laboratory) in 1% BSA in PBS. Finally, sections were washed in PBS, incubated with 4,6-diamidino-2-phenylindole (DAPI; Sigma 1μg ml^-1^), and mounted onto slides with Mounting Medium (Dako, #S3025)

### CLARITY

Full hemispheres were cleared based on a modified protocol^56^ derived from that described by Ye et al^57^. Briefly, >12 weeks old mice were transcardially perfused with ice cold PBS followed by 4% PFA, brains were removed and kept in 4% PFA overnight at 4#x00B0;C. Brains were then transferred to a 2% hydrogel solution (PBS with: 2% acrylamide, bio-rad #161-0140; 0.1% Bisacrylamide, bio-rad #161-0142; 0.25% VA-044 initiator, Wako, 011-19365; 4% PFA) for 48 hr. The samples were then degassed with N_2_ for 45 min and polymerized in 37#x00B0;C for 3.5 hr. After degassing, the samples were cut at the mid-sagittal plane. The samples were then washed overnight in 200 mM NaOH-Boric buffer (sigma, #B7901) containing 8% sodium dodecyl sulfate (SDS) (sigma, #L3771) to remove PFA residuals. Samples were then stirred in a clearing solution (100 mM Tris-Boric buffer, bio-lab, #002009239100 with 8% SDS) at 37#x00B0;C for 3–4 weeks. After the samples became transparent, they were washed with PBST (PBS with 0.5% tritonX100; ChemCruz, #sc-29112A) for 48 hr at 37#x00B0;C with mild shaking (replacing the PBST every 24 hr) and for another 24 hr with new PBST 0.5% at RT. Finally, the samples were incubated in the refractive index matched solution CLARITY Specific Rapiclear (RI = 1.45; SunJin lab, #RCCS002) O/N at 37#x00B0;C and for two more days at room temperature before imaging.

Transparent samples were embedded onto a slide surrounded by hot-glue walls. CLARITY specific RapiClear solution was then inserted into the chamber, covering the entire brain, and a thin coverslip glass was placed over the brain from above.

### Confocal Imaging

Confocal fluorescence images were acquired using an Olympus scanning laser microscope Fluoview FV1000 using 4X and 10X air objectives, 10X, and 20X water immersion objectives or 20X oil immersion objectives. Images were created by imaging between 50μm-5000μm in depth and reconstructed using IMARIS 9.1.2 software (Bitplane, UK).

### Imaris analysis

Using the ‘spot’ feature on the Imaris software, (x,y) coordinates were manually counted and extracted for all marked cells, DAPI included. To calculate the percentage of expected overlap between two groups out of a specific group, for each slice, the number of cells in the relevant group was divided by the total number of cells (i.e. DAPI) and multiplied by the same division for the second group in order to estimate the *percentage* of expected overlap. This number was then multiplied by DAPI in order to predict the *number* of estimated overlap cells, and later divided by the number of counted cells of the first group in order to estimate the percentage of overlap cells out of the relevant group. Finally, the percentage of actual overlap cells were calculated by dividing the empirical number of overlap cells by the number of counted cells from the first group. This parameter was later divided by the expected overlap, allowing us to calculate the *fold from expected* measurement:

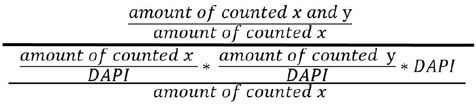

Where ‘*x’* represents the sub group of cells from which we wanted to extract the percentage (i.e. ‘how many of x cells were also y cells, out of the total amount of x cells’).

### Statistics

One-way Anova with Tukey or LSD post hoc and two tailed students t-test or paired t-test.

### Data Availability

The data used to support the conclusions of this study will be publicly available.

### Code Availability

Analysis codes will be made available to any interested reader.

## Acknowledgements

We thank the entire Goshen lab for their support. This project has received funding from the European Research Council (ERC) under the European Union’s Horizon 2020 research and innovation program (grant agreement No 803589), the Israel Science Foundation (ISF grants No. 1815/18 and 2060/23), and the Canada-Israel grants (CIHR-ISF, grant No. 2591/18). We thank Ami Citri and Nechama Novick for the critical reading of the manuscript.

This work is dedicated to the memory of Mrs. Lily Safra, a great supporter of brain research.

## References

1 Moscovitch, M., Cabeza, R., Winocur, G. & Nadel, L. Episodic Memory and Beyond: The Hippocampus and Neocortex in Transformation. Annual review of psychology 67, 105–134, doi:10.1146/annurev-psych-113011-143733 (2016).

2 Frankland, P. W. & Bontempi, B. The organization of recent and remote memories. Nat Rev Neurosci 6, 119–130, doi:10.1038/nrn1607 (2005).

3 Maviel, T., Durkin, T. P., Menzaghi, F. & Bontempi, B. Sites of neocortical reorganization critical for remote spatial memory. Science 305, 96–99, doi:10.1126/science.1098180 (2004).

4 Wiltgen, B. J., Brown, R. A., Talton, L. E. & Silva, A. J. New circuits for old memories: the role of the neocortex in consolidation. Neuron 44, 101–108, doi:10.1016/j.neuron.2004.09.015 (2004).

5 Frankland, P. W., Bontempi, B., Talton, L. E., Kaczmarek, L. & Silva, A. J. The involvement of the anterior cingulate cortex in remote contextual fear memory. Science 304, 881–883, doi:10.1126/science.1094804 (2004).

6 Bontempi, B., Laurent-Demir, C., Destrade, C. & Jaffard, R. Time-dependent reorganization of brain circuitry underlying long-term memory storage. Nature 400, 671–675, doi:10.1038/23270 (1999).

7 Einarsson, E. O. & Nader, K. Involvement of the anterior cingulate cortex in formation, consolidation, and reconsolidation of recent and remote contextual fear memory. Learn Mem 19, 449–452, doi:10.1101/lm.027227.112 (2012).

8 Gonzalez, C. et al. Medial prefrontal cortex is a crucial node of a rapid learning system that retrieves recent and remote memories. Neurobiology of learning and memory 103, 19–25, doi:10.1016/j.nlm.2013.04.006 (2013).

9 Ross, R. S. & Eichenbaum, H. Dynamics of hippocampal and cortical activation during consolidation of a nonspatial memory. The Journal of neuroscience : the official journal of the Society for Neuroscience 26, 4852–4859, doi:10.1523/JNEUROSCI.0659-06.2006 (2006).

10 Wiltgen, B. J. et al. The hippocampus plays a selective role in the retrieval of detailed contextual memories. Current biology : CB 20, 1336–1344, doi:10.1016/j.cub.2010.06.068 (2010).

11 Teixeira, C. M., Pomedli, S. R., Maei, H. R., Kee, N. & Frankland, P. W. Involvement of the anterior cingulate cortex in the expression of remote spatial memory. J Neurosci 26, 7555–7564, doi:10.1523/JNEUROSCI.1068-06.2006 (2006).

12 Fitzgerald, P. J. et al. Durable fear memories require PSD-95. Molecular psychiatry 20, 913, doi:10.1038/mp.2015.44 (2015).

13 Goshen, I. et al. Dynamics of retrieval strategies for remote memories. Cell 147, 678–689, doi:10.1016/j.cell.2011.09.033 (2011).

14 Vetere, G. et al. Chemogenetic Interrogation of a Brain-wide Fear Memory Network in Mice. Neuron 94, 363–374 e364, doi:10.1016/j.neuron.2017.03.037 (2017).

15 Wheeler, A. L. et al. Identification of a functional connectome for long-term fear memory in mice. PLoS Comput Biol 9, e1002853, doi:10.1371/journal.pcbi.1002853 (2013).

16 Araque, A., Parpura, V., Sanzgiri, R. P. & Haydon, P. G. Tripartite synapses: glia, the unacknowledged partner. Trends Neurosci 22, 208–215, doi:10.1016/s0166-2236(98)01349-6 (1999).

17 Haydon, P. G. GLIA: listening and talking to the synapse. Nat Rev Neurosci 2, 185–193, doi:10.1038/35058528 (2001).

18 Araque, A. et al. Gliotransmitters travel in time and space. Neuron 81, 728–739, doi:10.1016/j.neuron.2014.02.007 (2014).

19 Roth, B. L. DREADDs for Neuroscientists. Neuron 89, 683–694, doi:10.1016/j.neuron.2016.01.040 (2016).

20 Adamsky, A. et al. Astrocytic Activation Generates De Novo Neuronal Potentiation and Memory Enhancement. Cell 174, 59–71 e14, doi:10.1016/j.cell.2018.05.002 (2018).

21 Mederos, S. et al. Melanopsin for precise optogenetic activation of astrocyte-neuron networks. Glia 67, 915–934, doi:10.1002/glia.23580 (2019).

22 Suthard, R. L. et al. Chronic Gq activation of ventral hippocampal neurons and astrocytes differentially affects memory and behavior. Neurobiol Aging 125, 9–31, doi:10.1016/j.neurobiolaging.2023.01.007 (2023).

23 Iwai, Y. et al. Transient Astrocytic Gq Signaling Underlies Remote Memory Enhancement. Front Neural Circuits 15, 658343, doi:10.3389/fncir.2021.658343 (2021).

24 Wang, X. et al. Activation of astrocyte Gq pathway in hippocampal CA1 region attenuates anesthesia/surgery induced cognitive dysfunction in aged mice. Frontiers in aging neuroscience 14, 1040569, doi:10.3389/fnagi.2022.1040569 (2022).

25 Nagai, J. et al. Specific and behaviorally consequential astrocyte G(q) GPCR signaling attenuation in vivo with ibetaARK. Neuron 109, 2256–2274 e2259, doi:10.1016/j.neuron.2021.05.023 (2021).

26 Liu, S. et al. Astrocytes in CA1 modulate schema establishment in the hippocampal-cortical neuron network. BMC Biol 20, 250, doi:10.1186/s12915-022-01445-6 (2022).

27 Kol, A. et al. Astrocytes contribute to remote memory formation by modulating hippocampal-cortical communication during learning. Nat Neurosci 23, 1229–1239, doi:10.1038/s41593-020-0679-6 (2020).

28 Jones, M. E., Paniccia, J. E., Lebonville, C. L., Reissner, K. J. & Lysle, D. T. Chemogenetic Manipulation of Dorsal Hippocampal Astrocytes Protects Against the Development of Stress-enhanced Fear Learning. Neuroscience 388, 45–56, doi:10.1016/j.neuroscience.2018.07.015 (2018).

29 Yang, Q. et al. Optogenetic stimulation of CA3 pyramidal neurons restores synaptic deficits to improve spatial short-term memory in APP/PS1 mice. Progress in neurobiology 209, 102209, doi:10.1016/j.pneurobio.2021.102209 (2022).

30 Kol, A. & Goshen, I. The memory orchestra: the role of astrocytes and oligodendrocytes in parallel to neurons. Curr Opin Neurobiol 67, 131–137, doi:10.1016/j.conb.2020.10.022 (2021).

31 Lee, S. H., Mak, A. & Verheijen, M. H. G. Comparative assessment of the effects of DREADDs and endogenously expressed GPCRs in hippocampal astrocytes on synaptic activity and memory. Front Cell Neurosci 17, 1159756, doi:10.3389/fncel.2023.1159756 (2023).

32 Josselyn, S. A. & Tonegawa, S. Memory engrams: Recalling the past and imagining the future. Science 367, doi:10.1126/science.aaw4325 (2020).

33 Josselyn, S. A., Kohler, S. & Frankland, P. W. Finding the engram. Nature reviews. Neuroscience 16, 521–534, doi:10.1038/nrn4000 (2015).

34 Tonegawa, S., Pignatelli, M., Roy, D. S. & Ryan, T. J. Memory engram storage and retrieval. Current opinion in neurobiology 35, 101–109, doi:10.1016/j.conb.2015.07.009 (2015).

35 Silva, A. J., Zhou, Y., Rogerson, T., Shobe, J. & Balaji, J. Molecular and cellular approaches to memory allocation in neural circuits. Science 326, 391–395, doi:10.1126/science.1174519 (2009).

36 Refaeli, R., Kreisel, T., Groysman, M., Adamsky, A. & Goshen, I. Engram stability and maturation during systems consolidation. Curr Biol, doi:10.1016/j.cub.2023.07.042 (2023).

37 Martin, R., Bajo-Graneras, R., Moratalla, R., Perea, G. & Araque, A. Circuit-specific signaling in astrocyte-neuron networks in basal ganglia pathways. Science 349, 730–734, doi:10.1126/science.aaa7945 (2015).

38 Martin-Fernandez, M. et al. Synapse-specific astrocyte gating of amygdala-related behavior. Nat Neurosci 20, 1540–1548, doi:10.1038/nn.4649 (2017).

39 DeNardo, L. & Luo, L. Genetic strategies to access activated neurons. Curr Opin Neurobiol 45, 121–129, doi:10.1016/j.conb.2017.05.014 (2017).

40 DeNardo, L. A. et al. Temporal evolution of cortical ensembles promoting remote memory retrieval. Nat Neurosci 22, 460–469, doi:10.1038/s41593-018-0318-7 (2019).

41 Shpokayte, M. et al. Hippocampal cells multiplex positive and negative engrams. bioRxiv (2020).

42 Ye, L. et al. Wiring and Molecular Features of Prefrontal Ensembles Representing Distinct Experiences. Cell 165, 1776–1788, doi:10.1016/j.cell.2016.05.010 (2016).

43 Chung, K. & Deisseroth, K. CLARITY for mapping the nervous system. Nat Methods 10, 508–513, doi:10.1038/nmeth.2481 (2013).

44 Willard, A. M. & Gittis, A. H. Mapping neural circuits with CLARITY. Elife 4, e11409, doi:10.7554/eLife.11409 (2015).

45 Rogerson, T. et al. Synaptic tagging during memory allocation. Nature reviews. Neuroscience 15, 157–169, doi:10.1038/nrn3667 (2014).

46 Refaeli, R. et al. Features of hippocampal astrocytic domains and their spatial relation to excitatory and inhibitory neurons. Glia 69, 2378–2390, doi:10.1002/glia.24044 (2021).

47 Perea, G., Navarrete, M. & Araque, A. Tripartite synapses: astrocytes process and control synaptic information. Trends Neurosci 32, 421–431, doi:10.1016/j.tins.2009.05.001 (2009).

48 Nagai, J. et al. Behaviorally consequential astrocytic regulation of neural circuits. Neuron 109, 576–596, doi:10.1016/j.neuron.2020.12.008 (2021).

49 Durkee, C., Kofuji, P., Navarrete, M. & Araque, A. Astrocyte and neuron cooperation in long-term depression. Trends in neurosciences 44, 837–848, doi:10.1016/j.tins.2021.07.004 (2021).

50 Oliveira, J. F. & Araque, A. Astrocyte regulation of neural circuit activity and network states. Glia 70, 1455–1466, doi:10.1002/glia.24178 (2022).

51 Yu, X., Nagai, J. & Khakh, B. S. Improved tools to study astrocytes. Nature reviews. Neuroscience 21, 121–138, doi:10.1038/s41583-020-0264-8 (2020).

52 Durkee, C. A. et al. Gi/o protein-coupled receptors inhibit neurons but activate astrocytes and stimulate gliotransmission. Glia 67, 1076–1093, doi:10.1002/glia.23589 (2019).

53 Vaidyanathan, T. V., Collard, M., Yokoyama, S., Reitman, M. E. & Poskanzer, K. E. Cortical astrocytes independently regulate sleep depth and duration via separate GPCR pathways. eLife 10, doi:10.7554/eLife.63329 (2021).

54 Van Den Herrewegen, Y. et al. Side-by-side comparison of the effects of Gq- and Gi-DREADD-mediated astrocyte modulation on intracellular calcium dynamics and synaptic plasticity in the hippocampal CA1. Mol Brain 14, 144, doi:10.1186/s13041-021-00856-w (2021).

55 Madisen, L. et al. A robust and high-throughput Cre reporting and characterization system for the whole mouse brain. Nat Neurosci 13, 133–140, doi:10.1038/nn.2467 (2010).

56 Refaeli, R. & Goshen, I. Investigation of Spatial Interaction Between Astrocytes and Neurons in Cleared Brains. J Vis Exp, doi:10.3791/63679 (2022).

57 Ye, L. et al. Wiring and Molecular Features of Prefrontal Ensembles Representing Distinct Experiences. Cell 165, 1776–1788, doi:10.1016/j.cell.2016.05.010 (2016).

